# IgG sequencing maps systemic antibody binding to oral commensals and pathobionts in health and periodontitis

**DOI:** 10.64898/2026.09.28.753837

**Authors:** Ramin Akhi, Drake Winslow Williams, Teresa Greenwell-Wild, Andrew S Burns, Shreni Mistry, Laurie Brenchley, Nicolas Bouladoux, Ivan Vujkovic-Cvijin, Yasmine Belkaid, Akintunde Emiola, Niki M Moutsopoulos

## Abstract

The oral cavity harbors one of the body’s most diverse microbial communities, which continuously seeds the bloodstream and gastrointestinal tract, particularly in periodontitis, a disease in which both barrier permeability and microbial burden increase. Despite this constant systemic exposure, overt bacteremia from oral organisms is rare, suggesting an active surveillance system that contains this reservoir. Here, we apply IgG-sequencing to comprehensively map mucosal and systemic antibody recognition of the oral microbiome across paired oral fluid and serum from healthy individuals and patients with periodontitis. We find that IgG-binding is only weakly predicted by microbial abundance, suggesting taxon-specific and largely conserved IgG-binding of commensals, many of them streptococci which dominate antibody recognition irrespective of disease state. Periodontitis broadens the repertoire of IgG-bound taxa to include particular disease-associated pathobionts while preserving robust targeting of health-associated symbionts. Comparison of salivary and serum antibody profiles further uncovers a subset of commensal streptococci, including species implicated in infective endocarditis, that acquire disproportionate systemic IgG binding in periodontitis. Together, these findings establish a taxon-level atlas of humoral immunosurveillance at a major mucosal-microbial interface and reveal how periodontal disease reshapes systemic antibody recognition of both pathobionts and commensals.

## Introduction

The oral cavity is home to one of the richest and most diverse microbial communities in the human body, comprising several hundred bacterial taxa colonizing areas of the dentition, gingival crevice, tongue, and mucosal surfaces (Baker et al., 2024; Consortium, 2012). These communities, however, do not remain confined to their anatomical niche (Hajishengallis and Chavakis, 2021). Oral microbial translocation into the bloodstream occurs continuously, even during routine activities such as mastication and oral hygiene, and oral microbiota are further continuously disseminated via swallowed saliva into the gastrointestinal tract (Cobe, 1954; Lockhart et al., 2008; Schmidt et al., 2019). In health, this translocation is low-grade and has been thought of as immunologically inconsequential. In periodontitis however, where oral microbial burden increases logarithmically (Abusleme et al., 2013; Hajishengallis et al., 2023), and the epithelial barrier becomes increasingly permeable as a consequence of inflammation-driven tissue pathology (Moutsopoulos and Konkel, 2018; Theofilou et al., 2026), the potential for both hematogenous and alimentary microbial translocation rises substantially, positioning the periodontal pocket as a persistent portal through which oral microbiota can gain systemic access.

This continuous translocation of oral microbiota, which is amplified in periodontitis, has been implicated in the triggering and/or potentiation of distal and systemic disease manifestations. Periodontal pathogens have been linked epidemiologically and experimentally to cardiovascular disease, adverse pregnancy outcomes, rheumatoid arthritis, and glycemic dysregulation in diabetes, among other conditions (Brewer et al., 2023; Hajishengallis, 2015; Tonelli et al., 2023). These clinical associations and mechanistic insights reveal the potential of the oral cavity to serve as a systemic microbial reservoir with the capacity to influence distal organ physiology, particularly in the setting of periodontal disease.

Yet this reservoir is, for the most part, remarkably well contained. Episodes of clinically significant systemic bacteremia attributable to oral commensals or periodontal pathobionts remain rare, with the notable exception of infective endocarditis occurring in individuals with predisposing cardiac susceptibility (Chamat-Hedemand et al., 2020; Li et al., 2024). This low incidence of overt systemic morbidity suggests an active systemic surveillance mechanism directed against oral microbiota, one capable of containing translocating organisms before they establish clinically apparent disease elsewhere in the body.

Systemic antibody responses to oral microorganisms are detectable even in oral health, with salivary IgG derived predominantly from serum transudate and increasing in magnitude with periodontitis (Ogawa et al., 1989). Foundational work established that healthy individuals mount measurable systemic immune responses to their resident oral flora, indicating that this microbial population is continuously monitored rather than immunologically ignored. In periodontitis, increased antibody titers and specific recognition of oral pathobionts supports enhanced surveillance in the setting of disease (Ebersole et al., 1986). Perhaps the clearest illustration of microbe-specific systemic surveillance comes from a particular form of periodontal disease (often encountered in younger patients), characterized by hyper-elevated IgG2 responses directed specifically against *Aggregatibacter actinomycetemcomitans*, demonstrating that this humoral response can be both robust and remarkably pathogen-specific (Wilson et al., 1995).

Despite this evidence, a comprehensive, unbiased assessment of systemic antibody surveillance across the full breadth of the oral microbiota has been lacking. Therefore, we aimed to compile a comprehensive evaluation of IgG-binding of oral microbiota, with the intent to provide insights into the immune surveillance of oral taxa in health and periodontitis. We leveraged recent advances in IgG-sequencing technologies (Vujkovic-Cvijin et al., 2022) which allow for the characterization of antibody binding across the full diversity of the oral microbiome with a breadth and resolution not previously attainable and map the landscape of systemic IgG binding of this barrier’s resident microbial community in health and periodontal disease.

## Results

### Oral microbial communities reflect periodontal health status

To investigate antibody responses to oral microbiota in health and periodontitis, we recruited healthy participants and patients with periodontitis based on strict inclusion and exclusion criteria. Participants were screened for systemic health to exclude comorbidities and were categorized as healthy or severe/untreated periodontitis based on oral/periodontal evaluations (Suppl. Table 1). Research sampling included collection of gingival crevicular fluid (GCF, the local fluid at the site of the periodontal pocket adjacent to the tooth), saliva, and serum from each subject which allowed for studies of microbiome characterization and antibody binding at the tooth-associated area (GCF) and in the oral cavity overall (saliva) (Figure 1A).

**Figure 1.**
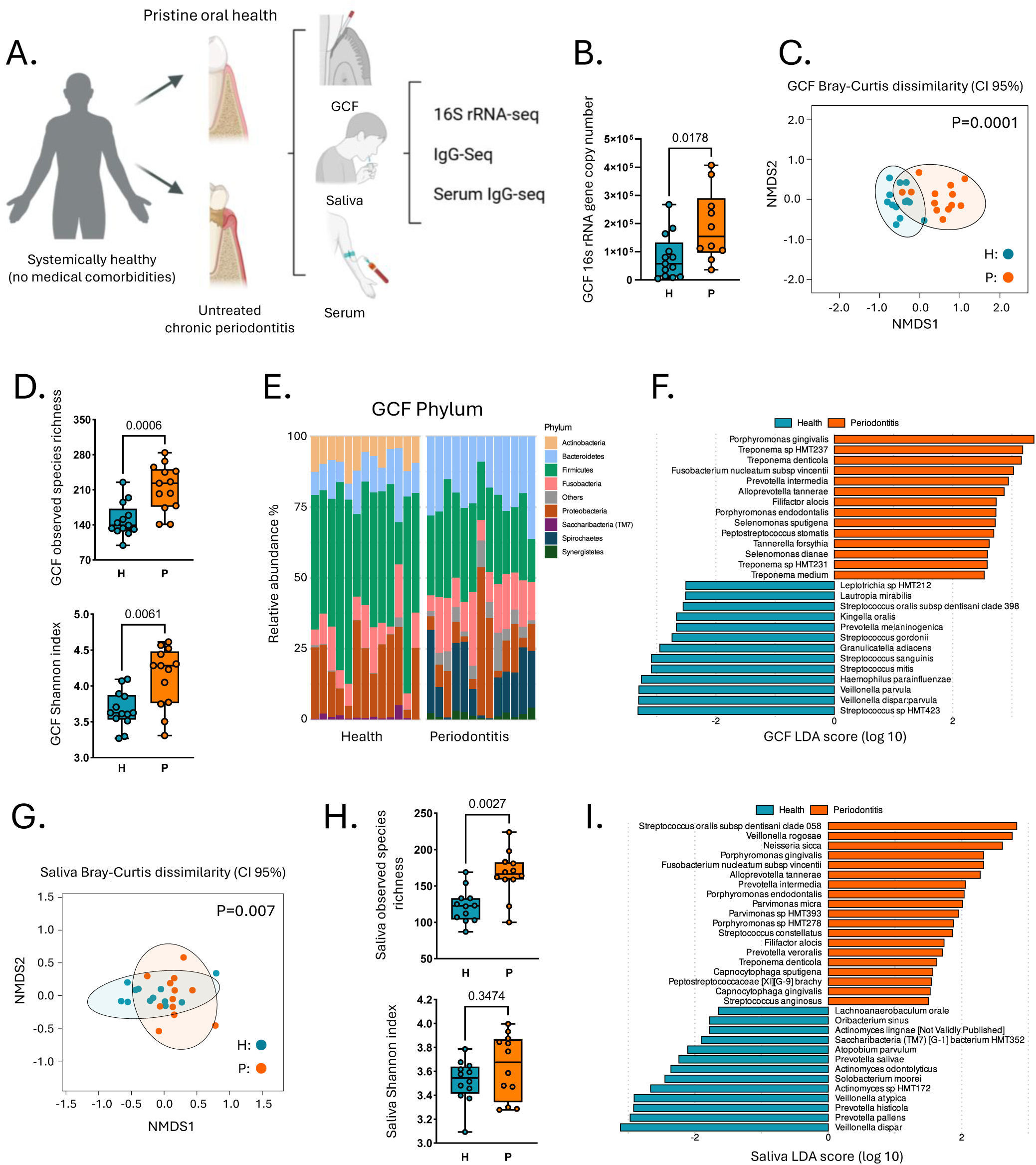
Cohort design and oral microbiome characterization. (A) Schematic of participant recruitment, clinical classification, and sample collection. (B) 16S rRNA gene copy number in gingival crevicular fluid (GCF) from healthy and periodontitis subjects. (C) Non-metric multidimensional scaling (NMDS) plot of GCF microbiome profiles from healthy (Cyan) and periodontitis (orange) samples based on Bray-Curtis dissimilarity test. (D) Observed species richness and Shannon index as measures of alpha diversity in GCF samples. (E) Taxonomic composition of GCF samples. Stacked bar charts showing relative abundance (%) of major bacterial phyla per sample. (F) Differentially abundant taxa in GCF between health and periodontitis identified by LEfSe (p < 0.05; LDA score> 2.5). (G) NMDS plot of salivary microbiome profiles from healthy (Cyan) and periodontitis (orange) samples based on Bray-Curtis dissimilarity analysis. (H) Observed species richness and Shannon index as measures of alpha diversity in saliva samples. (I) Differentially abundant taxa in saliva identified by LEfSe (p < 0.05; LDA score > 1.5). Panel A was created in BioRender. Akhi, R. For panels (B, D and H) group differences were assessed with two-sided Mann-Whitney U tests. In the NMDS plot, each dot represents one independent sample, and ellipses indicate the 95% confidence interval for each group. Group differences in community composition were assessed by PERMANOVA. For LEfSe, only taxa detected in at least 50% of subjects in either group, with a mean relative abundance >0.01% were included. Box plots indicate the median, interquartile range, and full data range. H: Healthy, P: Periodontitis.

Quantification of bacterial 16S rRNA gene copy number in tooth-associated GCF samples by qPCR revealed increased microbial biomass in periodontitis (Figure 1B) and distinct microbiome communities in periodontitis versus health (Figure 1C), as previously described (Abusleme et al., 2013). Periodontitis samples additionally exhibited significantly higher species richness and a higher Shannon Diversity Index (Figure 1D) (Iniesta et al., 2023; Ximénez-Fyvie et al., 2000). Profound differences between the health and periodontitis-associated microbial communities were also evident at the genus and phylum levels with periodontitis showing reduced abundance of *Actinobacteria* and expansion in *Bacteroidetes* and *Spirochaetes* phyla (Abusleme et al., 2013) (Figure 1E, Suppl. Figure 1A). Differential abundance analysis of microbial taxa also reflected significant differences between health and disease (Figure 1F) and included well-known periodontal pathobionts significantly enriched in the periodontitis samples, such as *Porphyromonas gingivalis, Treponema denticola, Prevotella intermedia, Filifactor alocis, Peptostreptococcus stomatis, Tannerella forsythia*, and *Selenomonas sputigena.* Conversely, microbes associated with oral health were underrepresented in the disease group (Figure 1F). This clear separation between health and periodontitis-associated microbial communities, was consistent with well-characterized features of oral microbiomes in health and disease and reflected the meticulous curation of our clinical cohorts and the accuracy of our sampling methods.

Matched saliva microbiome samples also displayed a distinct microbial composition with increased richness and beta-diversity in periodontitis compared to health (Figure 1G-H, Suppl. Figure 1B). LEfSe analysis of salivary microbiomes displayed differential abundance of taxa between health and disease, with over-representation of periodontitis-associated species in periodontitis saliva (Figure 1I). Interestingly, the relative abundance of detected taxa correlated significantly between matched saliva and GCF samples in both health and periodontitis (Suppl. Table 2), suggesting that salivary microbiomes mirror changes occurring at the area proximal to the tooth surface (Jung et al., 2024). In summary, our patient cohorts separate clearly based on health and periodontitis - associated microbiome signatures in both GCF and salivary oral fluids.

### IgG is present in oral fluids in health and increased in periodontitis

To characterize IgG-binding to oral microbiota we first measured total IgG across oral and systemic compartments. Salivary IgG was present at low concentrations in periodontally healthy subjects and was significantly elevated in patients with periodontitis. IgG in the tooth-associated fluid (GCF) also followed a similar trend (Figure 2A, Suppl. Figure 2A). Notably, GCF IgG concentrations correlated with disease severity, as measured by probing pocket depth (PPD), but not with microbiome biomass (Suppl. Figures 2B-C). As GCF contains IgG from serum transudate coming from the gingival tissue (Barros et al., 2016) and serum levels of IgG were not significantly different in health vs periodontitis (Figure 2B), we surmised that increased IgG GCF levels may be due to local antibody production within mucosal tissues. IgG positive cells have been previously documented in the mucosal tissues of patients with periodontitis (Ogawa et al., 1989; Williams et al., 2021) and the presence of tertiary lymphoid structures with the potential to support local antibody production has also been established in the lesions of severe periodontitis (Theofilou et al., 2026). Through re-analysis of existing single-cell datasets of human tooth - associated mucosa (gingiva) from healthy volunteers and patients with periodontitis we confirm the presence of few IgG-expressing plasma cells within tooth-associated mucosa (gingiva) in health, with a significant expansion in their numbers in periodontitis (Figure 2C) (Williams et al., 2021). Furthermore, immunostaining of gingival tissues from periodontitis patients in our cohort supports the presence of IgG positive plasma cells within oral mucosal lesions of disease (Figure 2D). Collectively, these data confirm that IgG, potentially derived from both serum and local production, is present in oral fluids and increased in patients with periodontitis.

**Figure 2:**
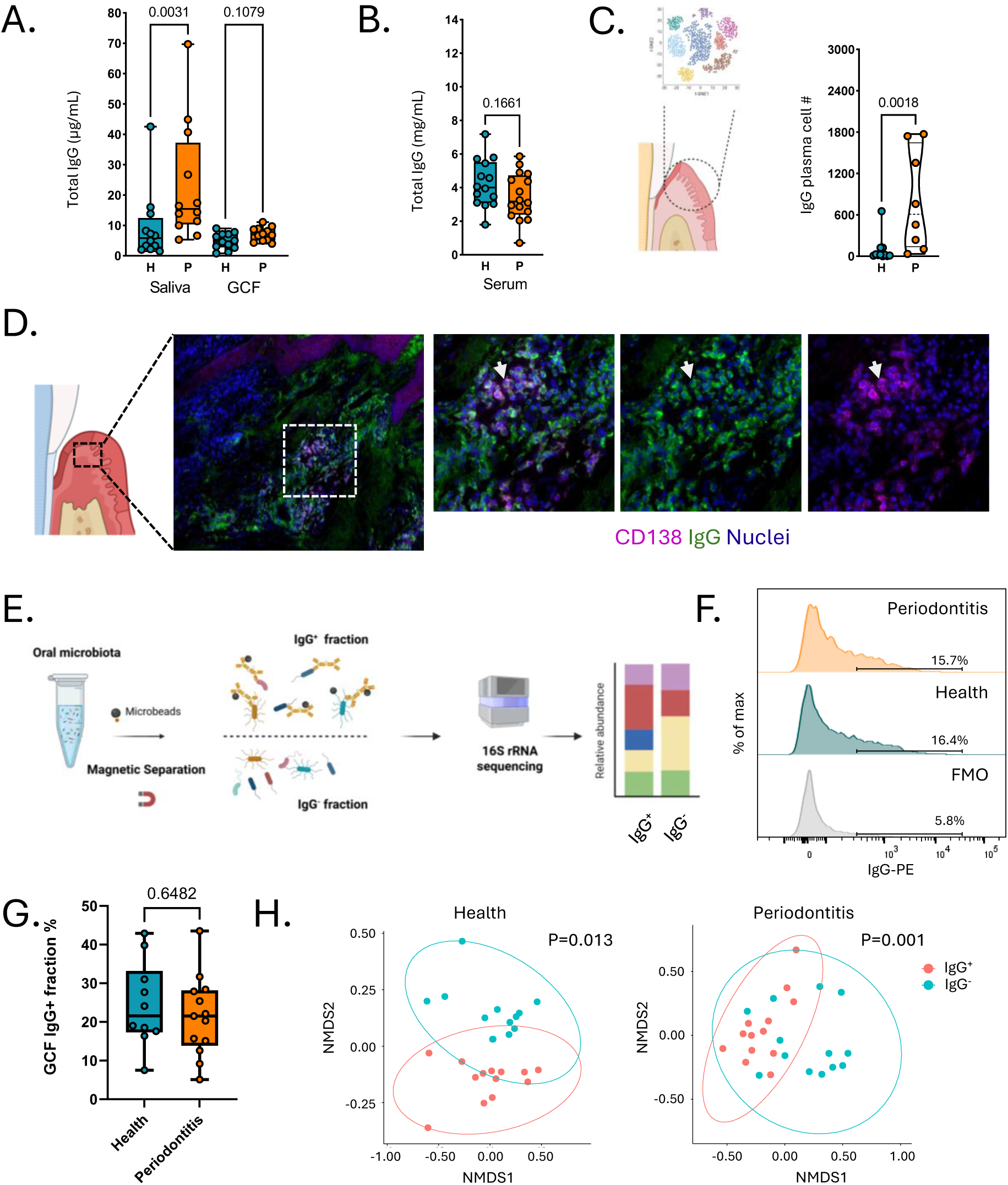
Humoral immunity and IgG sequencing methodology. (A) Total IgG levels in saliva, gingival crevicular fluid (GCF), and (B) serum across groups. Group differences were analyzed using two-sided Mann-Whitney U tests and Kruskal-Wallis tests, with p-values adjusted using the Benjamini-Hochberg (BH) method. (C) Cartoons illustrate gingival tissue sampling and single-cell RNA-seq analysis. IgG plasma cell counts per patient are shown on the right, grouped by disease status. Each dot represents one independent patient, and violin plots indicate the distribution within each group. (D) Schematic diagram of the gingival tissue indicating the region of interest (black dashed box). Representative immunofluorescence image of gingival tissue from a periodontitis patient, stained for CD138 (magenta), IgG (green), and nuclei (Hoechst, blue). The white dashed box indicates the region shown at higher magnification in panels. (E) IgG-seq workflow. An aliquot of oral microbiome sample was incubated with anti-IgG antibodies for flow cytometry analysis. IgG-coated bacteria were captured with microbeads, allowing magnetic separation of IgG-coated (IgG⁺) and IgG-uncoated (IgG⁻) fractions. Each fraction was subjected to 16S rRNA gene amplicon sequencing. (F) Flow cytometry analysis of IgG-coated oral bacteria. Representative histograms show PE-IgG signal in the fluorescence minus one (FMO) control and in health and periodontitis samples. The gate indicates the IgG-positive population and percentage denotes the fraction of IgG-coated bacteria in each sample. Histograms are shown as normalized event frequency. (G) Percentage of IgG-coated oral bacteria in health and periodontitis samples. Group differences were assessed using a two-sided Mann-Whitney U test. (H) NMDS plots of microbiome profiles in IgG⁺ and IgG^-^fractions from healthy and periodontitis samples based on Bray-Curtis dissimilarity calculated from 16S rRNA gene sequencing data. Panels C (left), D (left) and E were created in BioRender. Akhi, R. Each dot represents one independent sample, and ellipses denote the 95% confidence interval for each group. Group differences in community composition were assessed by PERMANOVA. Box plots show the median and interquartile range, with whiskers indicating the full data range; each dot represents one independent sample. H: Healthy, P: Periodontitis.

### IgG sequencing of tooth-associated microbial communities reveals distinct microbe binding in health and periodontitis

To profile IgG-binding to the oral microbiome, we employed an IgG-seq workflow that incorporates magnetic bead-based sorting (Vujkovic-Cvijin et al., 2022). Briefly, we labeled IgG-bound bacteria from oral fluids (GCF or saliva) with microbeads, enabling magnetic separation of IgG-coated (IgG^+^) and uncoated (IgG^-^) fractions for downstream 16S rRNA sequencing (Figure 2E). Flow cytometry analysis staining with PE-conjugated anti-IgG, revealed the proportion of antibody-bound bacteria in health and disease (Figure 2F-G). Ordination analysis from the sequencing data revealed separation of IgG^+^ and IgG^-^ bacterial fractions (Figure 2H), indicating that antibody-coated bacteria are compositionally distinct from uncoated bacteria.

We first aimed to further characterize IgG coating for oral taxa within the GCF and used sequencing-derived relative abundances to define an IgG Coating Index (ICI) for each taxon, calculated as the log_2_ enrichment of IgG^+^ over IgG^-^ (Figure 3A) (Palm et al., 2014). Taxa with an ICI > 4 were classified as highest coated (>94% IgG-coated), those with an ICI between 2 and 4 as high coated (80-94% IgG-coated), and those with an ICI <-4 as largely uncoated (>94% uncoated). Among the microbiota with highest and lowest ICI scores, in health *Streptococcus mitis* and *Streptococcus oralis* were highly abundant in the IgG^+^ fraction (>94% bound) while *Selenomonas noxia* was mostly unbound. In disease, the periodontal pathobionts *F. alocis* and *Parvimonas micra* were among the highest bound by IgG while *Selenomonas sputigena* was mostly unbound (Figure 3B).

**Figure 3.**
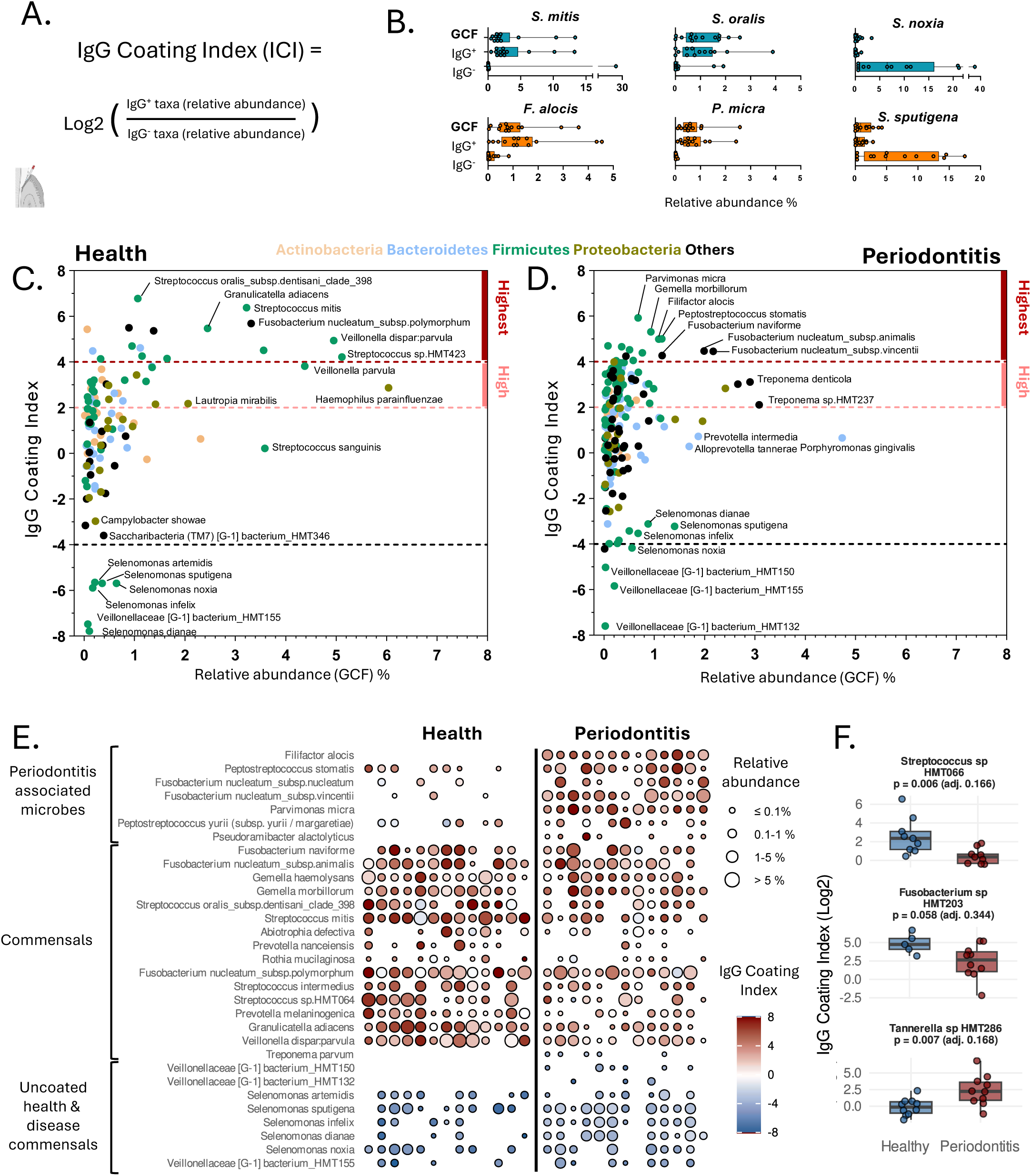
IgG-seq of the tooth-associated microbiome identifies differentially coated taxa in health and periodontitis. (A) For every taxon, an IgG Coating Index (ICI) value was calculated as the log2 ratio of its relative abundance in the coated versus the uncoated fractions. (B) Boxplots showing bacterial taxa with the highest and lowest IgG coating scores in healthy individuals and periodontitis patients. (C, D) IgG-coating profiles of gingival crevicular fluid (GCF) microbiota in healthy controls and participants with periodontitis. X-axis indicates mean relative abundance in the pre-enrichment fraction, and y-axis shows the ICI. Each point represents a taxon and is color-coded according to its bacterial phylum. Taxa with an ICI score between 4 and 8 (representing 94.1-99.6% coating; dark red on the right axis) are classified as highest coated, while those with an ICI score between 2 and 4 (representing 80.0-94.1% coating; light red line on right y axis) are considered high coated. In panels C and D, only taxa present in ≥50% of individuals in either group and with a mean relative abundance >0.01% are shown. (E) Heatmap of IgG-coating index and relative abundance for taxa highest coated and uncoated as identified in Figure 3C and 3D. Rows represent taxa, and columns correspond to individual participants. Color intensity denotes the ICI ranging from IgG-uncoated (blue) to highly IgG-highly coated (red). Circle size indicates the relative abundance of each taxon in the pre-enrichment fraction. (F) ICI for three bacterial taxa abundant in both groups, compared between healthy (blue) and periodontitis (red). Each point is one subject; boxes show the interquartile range and median, whiskers the min–max.; Wilcoxon rank-sum tests p-values unadjusted, with Benjamini-Hochberg adjusted values in parentheses. The gingival crevicular fluid sampling icon was created in BioRender. Akhi, R.

We next wanted to visualize the landscape of IgG binding in health and periodontitis and plotted each microbe’s mean ICI score against its mean relative abundance in the pre-enrichment fraction in either health or disease (Figure 3C-D). Taxa present in more than 50% of samples at an overall abundance >0.01% were displayed. Our first observation was that a larger number of taxa were high and highest bound in periodontitis compared to health (47 in health vs 74 in disease, Figure 3C-D), consistent with the increased richness of the periodontitis microbiome. A second finding was that there was only a modest correlation between a microbe’s abundance and its likelihood of being IgG-coated. While some abundant species were heavily targeted, many were not, and conversely, several low-abundance taxa were strongly coated by IgG (Suppl. Figure 3A). In healthy individuals, the highest IgG-coated bacteria included health-associated commensals such as *Streptococcus oralis, Streptococcus mitis, Granulicatella adiacens, and Rothia mucilaginosa*. In periodontitis, IgG strongly bound both key pathobionts (*F. alocis, Peptostreptococcus stomatis, P. micra*) but also health-associated symbionts (*S. oralis, S. mitis* and *G. adiacens*). Interestingly, the canonical periodontal pathogen *Porphyromonas gingivalis* was only partially IgG-coated in our dataset, while several *Selenomonas* and *Veillonella* species were consistently not bound with IgG in health and disease. Finally, phylum also did not appear to predict IgG-binding as IgG-coated taxa occurred across phyla.

To assess inter-individual variability in IgG-binding within groups, we created a heatmap visualizing both the relative abundance and ICI for the highest coated and uncoated taxa, derived from Fig. 3C-D for all patients in our cohort (Figure 3E). This analysis revealed highly consistent patterns within each group and provided an overview of taxa highest bound by IgG. Indeed, many of the high/highest bound taxa in the periodontitis microbiome were periodontal pathobionts significantly associated with disease (Figure 3E and Suppl. Figure 3B). However, many commensals *Streptococcus* and *Gemella* species were broadly coated across most individuals in both cohorts, though their abundance and ICI scores were typically higher in health. Taxa that were not IgG-bound were generally shared consistently across health and disease subjects (Figure 3E).

To further probe differences in antibody binding between health and periodontitis, we compared ICI values between groups for all species and displayed the 24 taxa with the lowest unadjusted p-values (Suppl. Figure 4A).We particularly focused on taxa in which differential relative abundance between health and disease could not explain differential binding (Suppl. Figure 4B) and discovered that a number of ubiquitous commensal taxa such as *Streptococcus sp. HMT066*, had higher ICI scores in health compared to disease (Figure 3F, Suppl. Figure 4B). Of particular interest *Fusobacterium sp. HMT203* was highest bound in health, despite a trend toward higher abundance, while *Tannerella sp. HMT286* displayed higher binding in periodontitis samples (Figure 3F, Suppl. Figure 4B).

Overall, this data provides an overview of IgG binding proximal to the tooth-associated biofilm (GCF) and reveals distinct patterns of antibody binding in health and disease.

### Salivary IgG-sequencing reflects binding of mucosal and tooth-associated microbiota

We next wanted to explore IgG-binding of microbiota in saliva as saliva contains a composite of microbes derived from all oral sites including tooth-associated and mucosal microbiomes (Jung et al., 2024). Given this property, we reasoned that saliva would be an ideal fluid for characterizing broad IgG recognition of the entire oral microbiome. We performed IgG-seq on saliva from our health and periodontitis cohorts, plotting IgG scores for all abundant taxa against their relative abundance within the unfractionated salivary microbiome, color coded according to phylum (Figure 4A-B). Highest bound taxa were both oral mucosal and tooth-associated commensals in both health and disease. As such, highest coated taxa (IgG score> 4) were predominantly *Firmicutes*, including multiple *Streptococcus* species, whereas *Selenomonas* species and *Campylobacter showae* were consistently uncoated, in both health and disease. Similar to the GCF, highly abundant taxa such as *Haemophilus parainfluenzae* and *Prevotella melaninogenica* showed comparatively low IgG coating, indicating that relative abundance of a taxon in saliva did not predict the degree of IgG binding. Additionally, assessment of inter-individual variability in salivary IgG coating (of highest coated/uncoated commensals) (Figure 4C), revealed overall consistent binding within groups.

**Figure 4.**
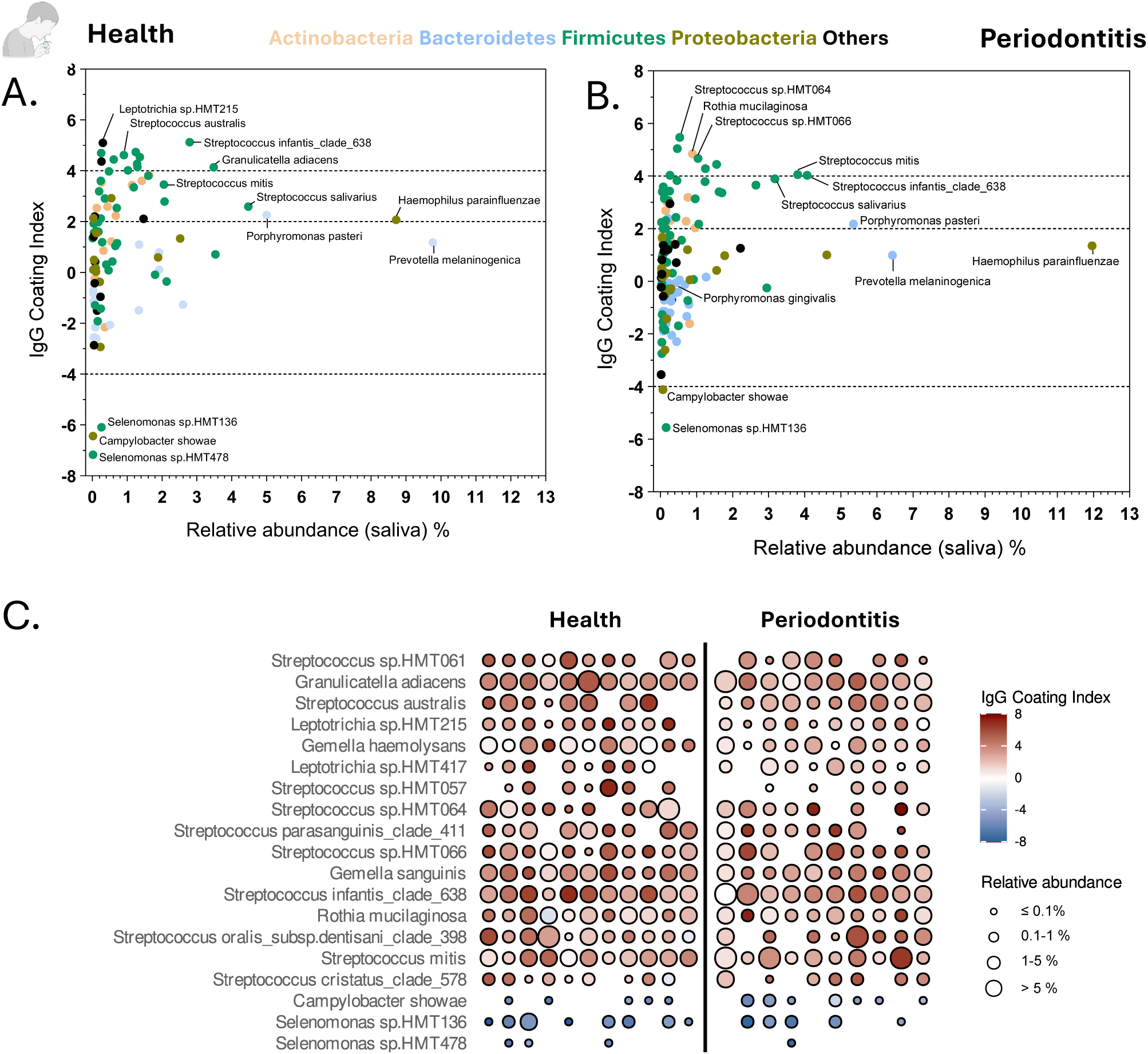
Salivary IgG-seq reveals oral microbiome coating profiles in health and periodontitis. (A, B) IgG-coating profiles of salivary microbiota color-coded according to its bacterial phylum in (A) healthy controls and participants with (B) periodontitis, x-axis shows the mean relative abundance and y-axis displays IgG Coating Index (ICI). Each point represents a taxon. (C) Salivary IgG-coating index and relative abundance shown as a heatmap for the most highly coated and uncoated taxa (Figure 4A and 4B). Each row is a taxon and each column an individual participant. The ICI is mapped by color, from uncoated (blue) to highly coated (red), while circle size reflects each taxon’s relative abundance in the pre-enrichment fraction. The saliva sampling icon was created in BioRender. Akhi, R.

Next, we investigated potential overlap in IgG-binding of microbial taxa between saliva and GCF in health and disease. For this purpose, we color coded taxa high and highest in GCF (derived from figure 3C-D), within the graphs displaying IgG-binding in saliva. Approximately half of the GCF high IgG-coated microbes (IgG score >2) were also highly coated in saliva samples (Suppl. Figure 5A-B, Suppl. Table 3). Various *Streptococcus* and *Gemella* species were highly coated in both cohorts, while in periodontitis, periodontitis-associated pathobionts such as *F. alocis* and *P. micra* were highly bound by IgG both in GCF and saliva (Suppl. Figure 5A, B). These results provide a comprehensive profile of IgG binding in saliva and suggest that saliva is a powerful sample for evaluating overall IgG-binding of oral microbiota.

### Serum IgG antibodies bind strongly to a subset of oral commensals

We further wanted to investigate IgG-binding to oral microbiota in the systemic circulation. Because saliva (and GCF) contains both serum and tissue derived IgG it is difficult to distinguish between systemic and locally derived IgG (Barros et al., 2016). We therefore reasoned that exploring the binding of oral microbiota by serum IgG can reveal taxa bound by circulating antibodies and that the comparison between serum and saliva IgG profiles would provide insights into taxa highest bound in the circulation. We thus incubated each participant’s salivary microbiome (which is already bound locally by IgG) with autologous serum and performed IgG-seq on the combined sample (Figure 5A). We calculated a serum ICI score to reflect the microbiota bound/unbound by the combination of saliva and serum IgG (Figure 5A). We plotted serum IgG scores against the relative abundance of each taxon (within saliva) in health and periodontitis. Serum IgG scores were generally higher (Figure 5B-C) than salivary IgG scores (Suppl. Table 4). Strikingly, the taxa with the highest serum IgG scores were mainly commensals belonging to the genus *Streptococcus* (including *S. australis, S. salivarius and S. HMT066*). Overall, there was significant overlap between IgG high/highest bound taxa in saliva and serum, but with additional high/highest bound taxa by circulating antibodies, particularly in periodontitis (Suppl. Figure 5C-D).

**Figure 5:**
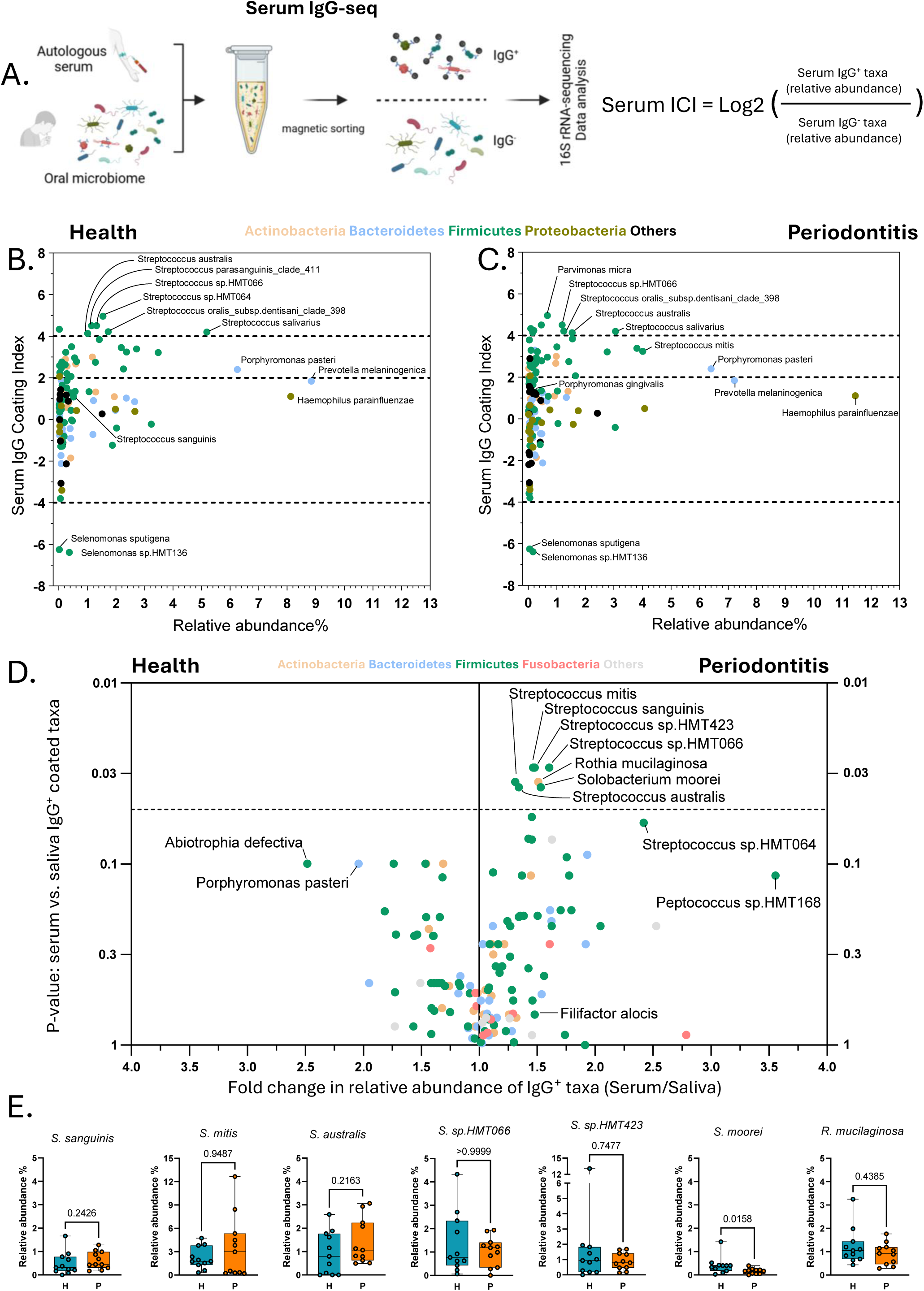
Serum IgG-seq maps systemic antibody targeting of oral commensals. (A) Schematic of the serum IgG-seq protocol. Salivary microbiota, already coated with salivary IgG, were incubated with autologous serum. IgG sequencing was then performed on the combined sample to assess serum IgG binding to oral microbes. (B, C) Serum IgG-coating profiles of oral microbiota in (B) healthy controls and individuals with (C) periodontitis. X-axis represents the mean relative abundance, and the y-axis indicates the ICI for each taxon. (D) Comparison of serum and salivary IgG coating of oral microbiota. The x-axis shows the fold change in relative abundance of IgG⁺-coated taxa (Serum/Saliva), and the y-axis displays adjusted p-values from statistical comparisons. Statistical significance was assessed using the Wilcoxon rank-sum test, and P-values were adjusted for multiple testing using the Benjamini-Hochberg method. Panels B-D are color-coded by bacterial phylum, and taxa detected in at least 50% of individuals in each group and with a mean relative abundance >0.01% are shown. (E) Relative abundance of species in saliva samples in healthy and periodontitis participants. Statistical significance was determined using the Mann-Whitney U test. Panel A was created in BioRender. Akhi, R.

To identify taxa with significantly higher binding by serum compared to salivary IgG, we compared per-taxon relative abundance in the serum IgG^+^ fraction versus the salivary IgG^+^ fraction. Intriguingly, a significant increase in serum IgG coating was found for a few commensal taxa including *Streptococcus mitis* and *Streptococcus sanguinis* in periodontitis but not in health (Figure 5D). This difference was not due to changes in relative abundance of these bacteria within the oral microbiome, as their abundance was not higher in periodontitis (Figure 5E). Notably, the species targeted by serum IgG in periodontitis (*i.e. S. mitis* and *S. sanguinis*) have been previously linked to systemic translocation and infective endocarditis in susceptible individuals (Li et al., 2024; Wilson et al., 2021). Collectively, these data suggest systemic recognition of specific oral commensals, including known opportunistic pathogens, in the setting of oral inflammatory disease.

## Discussion

Our study leverages IgG sequencing approaches to compile an atlas of IgG-binding to oral microbial taxa and provides an unbiased overview of IgG-binding to oral microbes in health and in periodontitis, a prevalent disease associated with microbial dysbiosis and systemic translocation of oral bacteria. The relevance of our findings may have broad implications towards dissecting host-microbiome immunity within the oral cavity but additionally, has the potential to open avenues towards understanding the participation of oral taxa in the induction of systemic immunity and the potentiation of distal disease. Indeed, oral microbiota are increasingly implicated in systemic conditions ranging from infective endocarditis to colon cancer, rheumatoid arthritis and Alzheimer’s disease (Dominy et al., 2019; Konig et al., 2016; Kostic et al., 2013).

Intriguingly, unlike other mucosal compartments (such as the gut), systemic IgG is ubiquitously present within the oral cavity, even in health, presumably modulating local barrier immunity in both health and disease. It has been established that serum IgG enters the oral cavity from the circulation through the permeable oral epithelial barrier (Hajishengallis, 2015; Ingrole et al., 2025). As a result, IgG is present within oral fluids, even in the setting of health (Akhi et al., 2017; Barros et al., 2016; Reinhardt et al., 1989). Yet, whether the totality of oral fluid IgG is serum-derived versus locally produced is hard to decipher, particularly in the context of oral inflammatory disease (periodontitis) in which tertiary lymphoid structures form within the mucosa and can support local production primarily of IgG (Easter et al., 2024; Ogawa et al., 1989; Sandholm and Grönblad, 1984; Theofilou et al., 2026; Williams et al., 2021). Importantly, although IgG binding to particular periodontal pathobionts is well established (Akhi et al., 2017; Schenck et al., 1987), a global overview of IgG binding to oral microbiota in health and periodontitis was lacking to date.

We performed IgG sequencing of oral fluids on a meticulously curated cohort of healthy participants and patients with severe, untreated periodontal disease. To minimize confounding variables, our inclusion criteria were strict, controlling for known systemic comorbidities, recent use of antibiotics, medications and other potential agents that can alter the host microbiome. These criteria, modeled after the original Human Microbiome Project sampling protocols (Dewhirst et al., 2010), were designed to ensure a highly controlled characterization of the microbiome and its antibody-binding profile in both health and periodontitis. Through this approach, we document broad IgG-binding to oral commensals within oral fluids in both health and periodontitis. In healthy participants, IgG-seq profiles revealed strong coating of commensal taxa such as *Streptococcus mitis*, *Streptococcus oralis*, and *Granulicatella adiacens* (Kolenbrander et al., 2010).

In periodontitis, a greater number of taxa were highly bound, reflecting increased diversity of microbiome and potentially an expanded antigenic repertoire. While ubiquitous commensals were bound in health and disease, in disease we additionally identify high IgG-coating of select known oral pathogens (Baker et al., 2024) such as *F. alocis*, *P. stomatis*, and *P. micra* suggesting an adaptive response to emergent oral pathobionts. This raises key questions about whether these microbes are particularly immunogenic and therefore elicit strong antibody responses and how antibody recognition affects colonization and virulence for these microbes. Answering these questions is critical for understanding how specific bacteria contribute to local and systemic inflammation. In contrast, the prototypic periodontal pathogen *P. gingivalis* did not show high antibody binding to IgG (Hajishengallis et al., 2011; Maekawa et al., 2014). Studies that move beyond IgG-seq to pinpoint specific antigenic targets and their physiological consequences are necessary to clarify how mucosal IgG immunity balances both commensals and pathobionts yet the current study can guide such mechanistic studies based on the unbiased profiling of IgG binding of the entire oral microbiome in health and disease.

A consistent finding across health and disease was the lack of antibody binding by certain microbes. Several *Selenomonas* species, emerging players in oral disease (Cho et al., 2023; Kim et al., 2024), remained almost entirely uncoated by IgG in both healthy and periodontitis samples. Such weak antibody recognition may confer a significant fitness advantage. By avoiding IgG binding, these bacteria may escape humoral immune constraints, such as aggregation, reduced *in situ* replication, and the enrichment of mucus-interacting functions (Olm et al., 2025). This immune invisibility could facilitate the persistence and pathogenic activity of *Selenomonas* within oral biofilms.

Beyond IgG-binding within oral fluids, we wanted to better characterize systemic IgG responses to oral bacteria in health and periodontitis within the systemic circulation. We therefore developed an approach that overcomes the limitations of using saliva alone, where IgG is diluted and includes locally produced antibodies originating from the mucosa (Ogawa et al., 1989). We incubated autologous serum and saliva from each individual and performed IgG-seq, allowing us to evaluate combined salivary and serum IgG binding. Intriguingly, the bacteria most highly bound by IgG in health and periodontitis were primarily oral commensals, not pathobionts which have elevated IgG responses in periodontitis. This finding may be clinically significant as the top IgG targets included species like *Streptococcus mitis* and *Streptococcus sanguinis,* which are associated with infective endocarditis in susceptible individuals (Lockhart et al., 2008; Wilson et al., 2021). Remarkably, despite the increased barrier permeability and inflammation in periodontitis, these patients do not have higher incidence of endocarditis (Lockhart et al., 2008). We suspect that these systemic IgG responses may be protective by serving to neutralize translocating commensals and limit systemic bacteremia in patients with periodontitis. This interpretation aligns with previous gut studies showing that anti-microbiota IgG marks barrier-crossing taxa and can both opsonize and protect against systemic infection (Vujkovic-Cvijin et al., 2022; Zeng et al., 2016).

Collectively, this work provides a foundational atlas of antibody binding to oral commensals and pathobionts, revealing key features of antibody-mediated immunity to oral microbes in health and oral inflammatory disease.

## Supplemental Figures/Tables legend

**Supplemental Table 1.** Demographic characteristics of study participants and experiments performed for each individual.

**Supplemental Table 2.** Subject-wise Spearman correlation between relative abundance of Saliva and Gingival Crevicular Fluid (GCF).

**Supplemental Table 3.** GCF and Salivary IgG ICI scores for taxa highly coated in GCF.

**Supplemental Table 4.** Salivary and serum ICI scores by taxon in health and periodontitis.

**Supplemental Figure 1. Genus-level microbial profiles of GCF and saliva in health and periodontitis.** (A and B) Stacked bar plots showing the relative abundance of bacterial genera in gingival crevicular fluid (GCF) (A) and saliva (B) from healthy individuals and patients with periodontitis. Each bar represents an individual participant.

**Supplemental Figure 2.** G**C**F **IgG levels are elevated in periodontitis and correlate with probing pocket depth but not bacterial load.** (A) GCF IgG fold increase in periodontitis relative to healthy controls (periodontitis/healthy). (B) Correlation between GCF IgG concentrations and periodontal disease severity, quantified as probing pocket depth (PPD, mm). (C) Correlation of GCF IgG concentrations with GCF bacterial burden, quantified by 16S rRNA gene copy number. Each dot represents one participant; r and p values are indicated.

**Supplemental Figure 3.** I**g**G **coating index in GCF.** (A) Correlation analysis between taxon relative abundance and IgG Coating Index (ICI) scores, using Spearman’s rank correlation with two-tailed P-values. (B) Heatmap displaying IgG Coating Index (ICI) of taxa highly abundant in healthy and periodontitis participants derived from figure 1F. Rows represent taxa, and columns correspond to individual participants. Color intensity denotes the ICI, ranging from IgG-uncoated (blue) to highly IgG-coated (red). Circle size indicates the relative abundance of each taxon.

**Supplemental Figure 4.** T**o**p **24 GCF taxa ranked by IgG-coating difference between groups.** (A) IgG Coating Index (log2) and (B) relative abundance (%) for the same 24 taxa, ordered by ascending P-value. Boxplots show median, interquartile range, and range, with individual data points overlaid. P-values (raw and Benjamini-Hochberg adjusted) from Wilcoxon rank-sum tests are shown above each panel. Healthy, blue; periodontitis, red

**Supplemental Figure 5**. S**a**livary **and serum IgG microbiome coating profiles.** (A, B) IgG-coating profiles of salivary microbiota in (A) healthy controls and participants with (B) periodontitis; the x-axis shows the mean relative abundance and the y-axis displays ICI. Taxa are color-coded based on their GCF IgG-coating status (Figure 3C-D): dark red indicates highest coated taxa in GCF (ICI 4-8), and light red indicates high coated taxa in GCF (ICI 2-4). (C, D) Venn diagrams comparing high and highest coated taxa (ICI ≥ 2) in the salivary microbiome at baseline (salivary IgG) versus after sequential addition of serum IgG, shown for (C) healthy controls and (D) periodontitis. The overlap indicates taxa coated by both salivary and serum IgG; the saliva-only region indicates taxa coated by salivary IgG alone at baseline; and the serum region indicates taxa coated fraction after serum IgG coating.

## Materials and methods

### Clinical characterization and human samples collection

Participants were enrolled by NIH Clinical Center IRB approved protocol (ClinicalTrials.gov # NCT01568697) and provided written informed consent. Human participants were screened with rigorously defined inclusion and exclusion criteria for systemic and oral health. Healthy participants had minimal gingival inflammation and normal bone support. Participants with periodontitis were all diagnosed with stage III/IV disease based on AAP/EFP classification (Caton et al., 2018) and had no history of periodontal treatment. Both healthy and periodontitis participants were non-smokers and had no known systemic health conditions that could otherwise influence immune responses and periodontal health (i.e. diabetes, autoimmune conditions, pregnancy) and no use of medications including antibiotics for more than 3 months.

### Saliva and GCF collection and processing

Unstimulated saliva was collected on ice into 50-mL conical tubes over a 5-min interval. Samples were transferred onto a 40-µm filter placed atop a second 50-mL tube and centrifuged at 500 × g for 2 min at 4 °C, the filter was then discarded. The flow-through was further centrifuged at 3,000 × g for 5 min at 4 °C to pellet cellular material. The supernatant was removed, and the pellet was resuspended in 400 µL of 15% (v/v) glycerol. Suspensions were stored at −80 °C until use. Tooth-associated areas were isolated with cotton rolls to avoid active saliva contamination and GCF was collected with Periopaper strips and placed directly into 400 µL of 15% (v/v) glycerol buffer. For processing, an additional 100 µL of the same buffer (4 × 25 µL) was applied to each strip, followed by centrifugation at 2,000 rpm for 5 min at 4 °C. Samples were subsequently centrifuged at 15,000 rpm for 5 min at 4 °C, 450 µL of supernatant was discarded, and the pellet was resuspended in the remaining 50 µL. Samples were stored at −80 °C until use.

### Blood collection and serum processing

Peripheral venous blood was collected from each subject by standard phlebotomy according to the existing clinical protocol. Samples were centrifuged at 1,800 × g for 20 min at room temperature. Serum was separated, aliquoted into screw-cap cryovials, and stored at −80 °C until analysis.

### IgG-seq on microbiome of oral fluids

To assess IgG coating of the oral microbiota, we employed a high-throughput plate-based IgG-seq approach (Vujkovic-Cvijin et al., 2022) with slight modifications. Briefly, saliva and gingival crevicular fluid (GCF) samples were normalized at an equivalent optical density at 600 nm (OD600). Each sample was divided into different aliquots: two designated for IgG-seq, and one native sample. Aliquots were added to a 96-well plate, pelleted (6000 × g for 2 minutes at 4°C), washed with 1X PBS containing 1% bovine serum albumin (BSA) buffer, and resuspended in 20 μl of immunoglobulin-specific magnetic microbeads. Microbeads were made by conjugating reduced recombinant Protein A/G or Protein G (Cat# NBP2-34999 and Cat# NBP2-34962, Novus Biologicals) to HiSur Amine Beads (Cat# HA0153, Ocean Nanotech) using LC-SMCC [succinimidyl 4-(N-maleimidomethyl) cyclohexane-1-carboxy-(6-amidocaproate)] (Cat# 22362, ThermoFisher) using the Ocean Nanotech manufacturer’s protocol. The bacteria-microbead suspension was shaken at 400 rpm for 20 min at room temperature and then allowed to separate by incubation on a 96-well plate magnet for 3 min. Both the bead fraction (IgG-coated; IgG^+^) and supernatant (IgG-uncoated; IgG^-^) were transferred to −80°C for subsequent DNA extraction.

DNA was extracted from IgG^+^, IgG^-^, fractions and native samples using the Qiagen MagAttract PowerMicrobiome DNA/RNA Isolation Kit on an Eppendorf automated liquid handling system as detailed by the manufacturer. The V1V2 region of the 16S rRNA gene was amplified using the 8F (5′-AGAGTTTGATCMTGGCTCAG-3′) and 431R (5′-CYNACTGCTGCCTCCCGTAG-3′) with sample-specific barcodes and Illumina sequencing adapters appended to the primers (Kozich et al., 2013). Amplicons were purified using AMPure XP beads (Beckman Coulter), quantified via the KAPA Library Quantification Kit (Roche), and pooled at equimolar concentrations. Pooled libraries were sequenced on the Illumina MiSeq instrument utilizing the 600 cycle v3 Paired-End (250×250) Reagent Kit.

Demultiplexed paired-end 16S rRNA gene sequencing reads were processed using the DADA2 (Callahan et al., 2016) workflow. Reads were quality filtered, trimmed, denoised, and merged to generate high-resolution ASV (Amplicon Sequence Variant). Taxonomy was assigned using Human Oral Microbiome Database (HOMD) (Chen et al., 2010) in the R programming environment to construct ASV table. ASV abundance tables were merged with taxonomic classifications and aggregated at the genus and species levels for downstream analyses. Relative abundance was calculated for each sample by normalizing taxon-specific read counts to the total number of reads per sample.

IgG coating index (ICI) were calculated as log2 (IgG^+^/IgG^-^) for each taxon. For taxon with detectable IgG^+^ relative abundance and a zero or missing IgG^-^ value, the denominator was imputed using the lowest non-zero abundance of the corresponding taxon within the relevant group. We calculated the percentage of IgG coating by converting log2-transformed ratios into linear values. To illustrate, an IgG^+^/IgG^-^ ratio of 4 on a log2 scale corresponds to a linear ratio of 2^4^=16, indicating that the IgG^+^ is 16-fold higher than the IgG^-^ (a 16:1 relationship). To determine the IgG^+^ fraction relative to the total population IgG^+^/(IgG^+^ + IgG^-^), we treated the IgG^-^ as one reference unit and the IgG^+^ as 16 units, resulting in a total of 17 units. Thus, the IgG^+^ fraction is calculated as 16/ (16+1) =0.941, or 94.1%. Similarly, a log2 ratio of 0 corresponds to a linear value of 1 (2^0^=1), representing an equal distribution between IgG^+^ and IgG^-^ (a 1:1 relationship), which yields a coating percentage of 50%. The differential abundance microbial taxa were analyzed using LEfSe (Linear Discriminant Analysis Effect Size) package (Segata et al., 2011). For LEfSe analysis and ICI panels, taxa with relative abundance below 0.01% of the total and detected in fewer than 50% of samples within each group were excluded. Beta diversity was assessed in R using the vegan package based on Bray-Curtis dissimilarity and visualized by non-metric multidimensional scaling (NMDS), whereas alpha diversity was calculated from sample-by-taxon abundance matrices. Data wrangling and visualization were performed in R using the dplyr, tidyr, RColorBrewer and ggplot2 packages. Plots also were generated in GraphPad Prism 9 (GraphPad Software).

### Quantification of total bacteria

The determination of the total bacterial load was performed via qPCR using 16S rRNA gene primers (Abusleme et al., 2013). Total genomic DNA from *P. gingivalis* ATCC 33277 was used to construct standard curves with a known number of 16S rRNA gene copies. The qPCR reactions had a final volume of 20 µL and included 10 µL of SYBR Green PCR Master Mix (Applied Biosystems 4309155), 2.5 µM of reverse and forward primer (final 250 nM) (Barman et al., 2008), 1 µL of DNA from each sample and PCR water (MoBio). Thermal cycle conditions were 50°C for 2 min, 95°C 2 min and 40 cycles of 95°C for 15 s and 60°C for 1 min. All PCR reactions were performed in duplicate and were carried out in a 7500 Real-Time PCR system (ThermoFisher Scientific).

### Sc-RNA data reanalysis

Plasma cells were identified based on prior annotations (Williams et al., 2021) and further classified into IgG-expressing subsets according to immunoglobulin heavy chain gene expression profiles. The disease status of individual patients was used to quantify and compare IgG plasma cell frequencies across individuals and clinical groups.

### Immunofluorescence Staining

Five-micrometer fixed-frozen gingival tissue sections obtained from three periodontitis patients were rehydrated in 1× PBS at room temperature for 5 minutes. Sections were then blocked in 0.22 μm-filtered 1× PBS containing 0.3% Triton-X-100 (VWR Life Sciences), 1% BSA (Cat. A1933-5G, Sigma), 1% human Fc block (Cat. 422301, Biolegend), and 0.02% Hoechst 33342 (Cat. 40046, Biotium) at 37°C for 1 hour. Primary antibodies were applied overnight at 4°C and included anti-CD138 conjugated to Alexa Fluor 647 (mouse IgG1, clone Syndecan-1; BioLegend, cat# 356523; 1:200) and anti-IgG conjugated to Alexa Fluor 488 (rabbit monoclonal; Abcam, cat. ab307524; 1:200). Subsequently, sections were washed with 1× PBS. Nuclei were counterstained with Hoechst included in the blocking buffer. The staining yielded consistent results across all three patients. Fluorescence images were acquired using a Nikon TiE widefield microscope and processed using ImageJ (National Institutes of Health, Bethesda, MD, USA).

### Flow cytometry

Oral samples were pelleted at 6,000 x g for 2 min at 4°C, washed with 1× PBS, and stained with LIVE/DEAD Fixable Near-IR Dead Cell Stain (Invitrogen) for 20 min at 4°C. Cells were washed with 1× PBS containing 1% BSA and stained with anti-human IgG-PE (Miltenyi Biotec, Cat# 130-119-878). Stained samples were acquired using an LSRII flow cytometer (BD Biosciences). Flow cytometry data were analyzed in FlowJo v10.10.0.

### Serum IgG-seq

The serum from participants was heat-inactivated at 56°C for 35 minutes to block complement activity and was diluted to 1:300. Aliquots for IgG fractionation were incubated with sera from the same individual for 30 minutes at 4°C before proceeding with the magnetic bead-based fractionation. The fractionation, staining, sequencing, and data analysis steps were performed as described above for the oral fluids IgG-seq method.

### Antibody measurements

Salivary, GCF and serum immunoglobulins were quantified by ELISA using human IgG (BMS2091; Thermo Fisher Scientific) kits according to the manufacturer’s instructions. IgG levels in the healthy and periodontitis groups were measured from GCF and unstimulated whole saliva samples. Given that GCF samples in healthy subjects were exhausted in IgG-seq experiments, a matched healthy cohort was used. For GCF elution, Periopaper strips from two sites were placed into 100 µL assay buffer in 1.5-mL microcentrifuge tubes and mixed by Thermomixer at 1,000 rpm for 15 min at room temperature; strips were then positioned across the inner rim of the tube cap and centrifuged at 10,000 × g for 5 min to elute absorbed fluid, rinsed with 50 µL assay buffer, and centrifuged again at 10,000 × g for 10 min. The resulting supernatants were collected for analysis. Because salivary and GCF antibody concentrations differ between health and periodontitis, a pilot serial-dilution experiment was used to establish working dilutions: salivary IgG 1:200 and GCF IgG 1:20. Immunoglobulin concentrations were calculated from standard curves, and between-group differences were assessed with the Kruskal-Wallis test; data were plotted in GraphPad Prism 9 (GraphPad Software).

### Statistical methods

Differences in beta diversity were tested by permutational multivariate analysis of variance (PERMANOVA). Differences between two groups were assessed using the Mann-Whitney U test, whereas comparisons among three or four groups were performed using the Kruskal-Wallis test. For analyses involving multiple pairwise comparisons, the Wilcoxon rank-sum test was used, with P values adjusted for multiple testing by the Benjamini-Hochberg method. Correlation analysis was performed using Spearman’s rank correlation test.

## Data availability

The raw 16S rRNA gene amplicon sequencing data generated in this study have been deposited in the NCBI Sequence Read Archive (SRA) under BioProject accession PRJNA1365893.

## Acknowledgments

This research was supported [in part] by the Intramural Research Program of the National Institutes of Health (NIH). The contributions of the NIH author(s) are considered Works of the United States Government. The findings and conclusions presented in this paper are those of the author(s) and do not necessarily reflect the views of the NIH or the U.S. Department of Health and Human Services. This work was supported by the Intramural Program, Division of Intramural Research of the National Institute of Allergy and Infectious Diseases (NIAID), the Intramural Program of the National Institute of Dental and Craniofacial Research (NIDCR), and NIDCR extramural #R00DE032747 (D.W.W.), National Heart, Lung, and Blood Institute extramural #DP1HL174182 (I.V.-C.), and by the Finnish Cultural Foundation (R.A.). We thank the Spanish Catholic Center of Catholic Charities Dental Services, Catholic Charities of the Archdiocese of Washington, and the Department of Health and Human Services Dental Service, Maryland County Government, for their collaboration and for assisting with the recruitment of some study participants.

## Author contributions

R.A., D.W.W., and N.M.M. conceived and designed the study. R.A., D.W.W., A.B., S.M., T.G.W., and N.B. performed experimental work. T.G.W. processed participants samples. L.B. was involved in clinical cohort recruitment, patient evaluation, and clinical and research sample collection. Y.B. consulted on study design. I.V.-C. and A.E. were consulted on data analysis. R.A. performed data analysis and data interpretation. R.A., D.W.W. and N.M.M. wrote the manuscript. N.M.M. contributed to data interpretation and supervised the study. All authors read and approved the manuscript.

