## Supplementary figures and images for "IgG sequencing maps systemic antibody binding to oral commensals and pathobionts in health and periodontitis"

### Suppl figure 1-4

A.

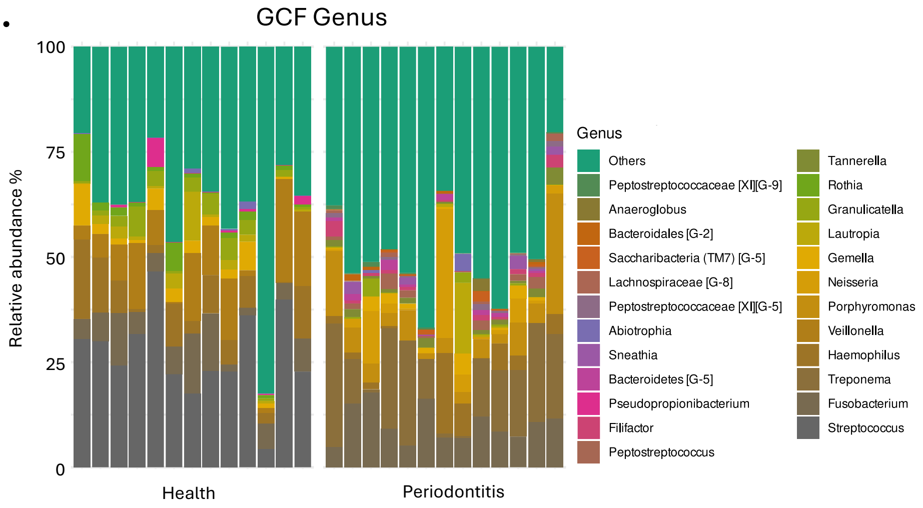


B.

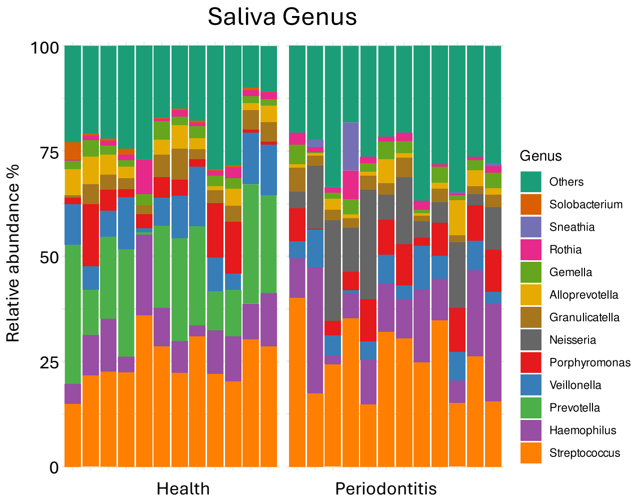


Suppl. Figure 1

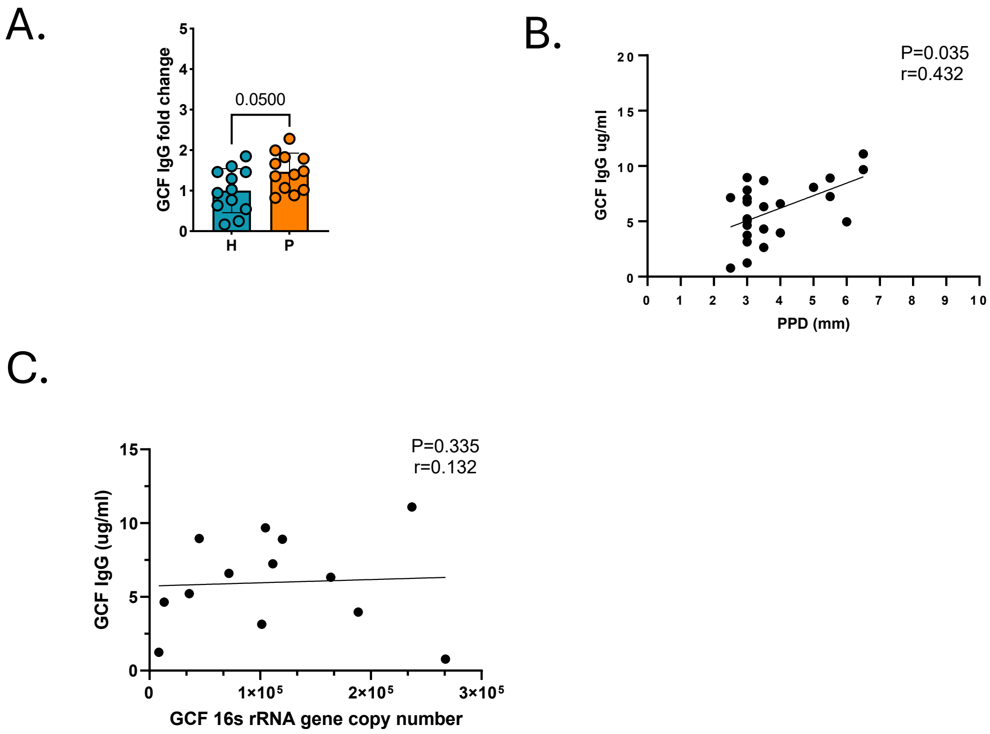


Suppl. Figure 2

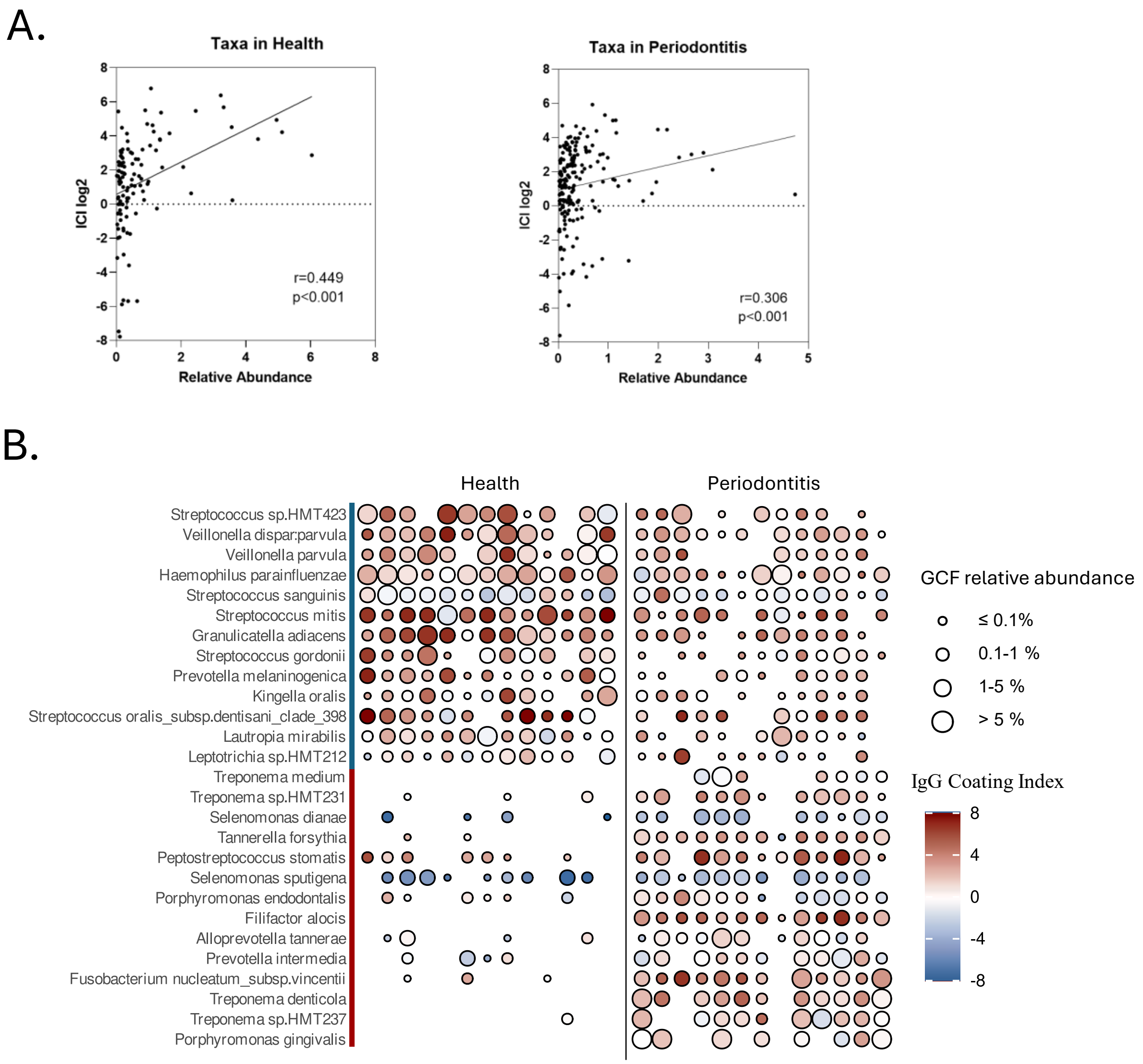


Suppl. Figure 3

A.

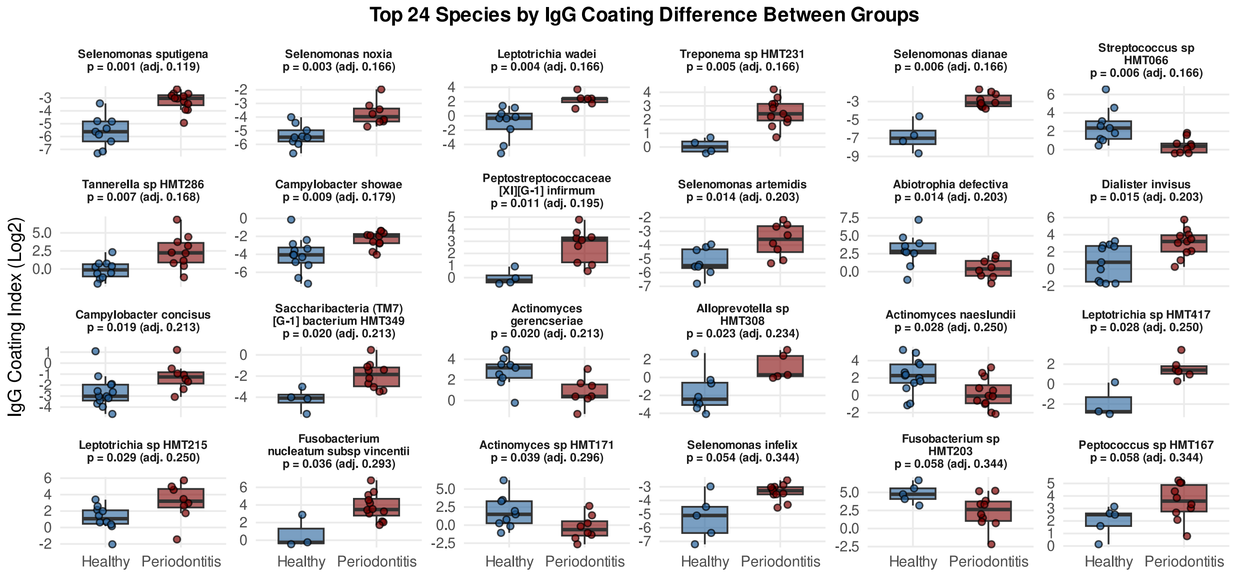


B.

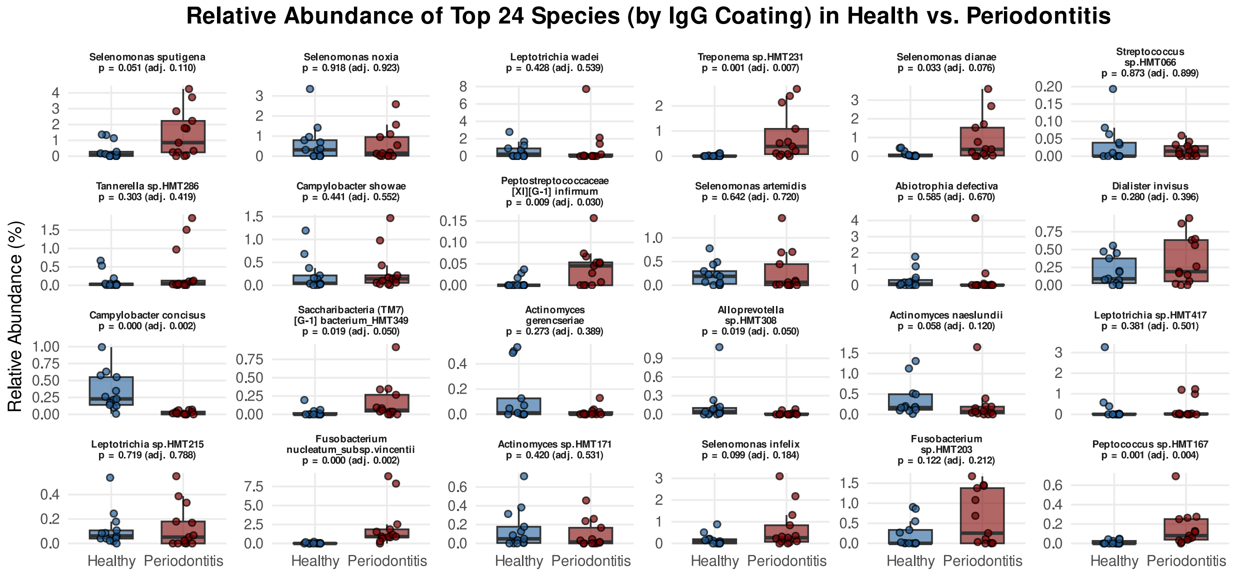


Suppl. Figure 4

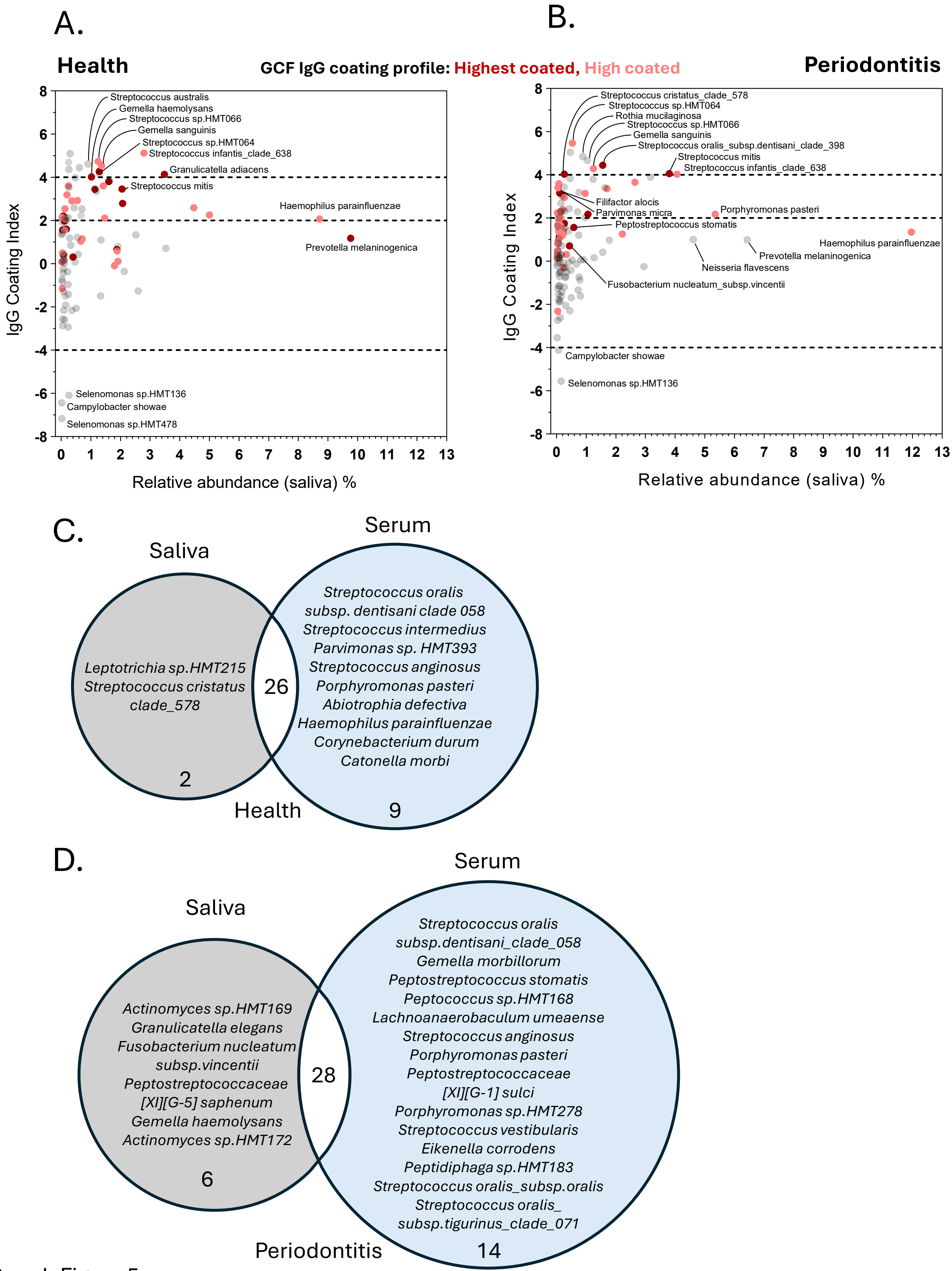


Suppl. Figure 5
